# mTOR inhibition *via* Rapamycin treatment partially reverts the deficit in energy metabolism caused by FH loss in RPE cells

**DOI:** 10.1101/2021.10.29.466270

**Authors:** David A. Merle, Francesca Provenzano, Mohamed Ali Jarboui, Ellen Kilger, Simon J. Clark, Michela Deleidi, Angela Armento, Marius Ueffing

## Abstract

Age-related macular degeneration (AMD) is a complex degenerative disease of the retina with multiple risk-modifying factors, including ageing, genetics and lifestyle choices. The combination of these factors leads to oxidative stress, inflammation and metabolic failure in the retinal pigment epithelium (RPE) with subsequent degeneration of photoreceptors in the retina. The alternative complement pathway is tightly linked to AMD. In particular, the genetic variant in the complement factor H gene (*CFH*), that leads to the Y402H polymorphism in the factor H protein (FH), confers the second highest risk for the development and progression of AMD. While the association between the FH Y402H variant and increased complement system activation is known, recent studies have uncovered novel FH functions not tied to this activity and highlighted functional relevance for intracellular FH. In our previous studies, we show that loss of *CFH* expression in RPE cells causes profound disturbances in cellular metabolism, increases the vulnerability towards oxidative stress and modulates the activation of pro-inflammatory signaling pathways, most importantly the NF-kB pathway. Here, we silenced *CFH* in hTERT-RPE1 cells to investigate the mechanism by which intracellular FH regulates RPE cell homeostasis. We found that silencing of *CFH* results in hyperactivation of mTOR signaling along with decreased mitochondrial respiration and that mTOR inhibition *via* rapamycin can partially rescue these metabolic defects. To obtain mechanistic insight into the function of intracellular FH in hTERT-RPE1 cells, we analyzed the interactome of FH via immunoprecipitation followed by Mass spectrometry-based analysis. We found that FH interacts with essential components of the ubiquitin-proteasomal pathway (UPS) as well as with factors associated with RB1/E2F signalling in a complement-pathway independent manner. Moreover, we found, that FH silencing affects mRNA levels of the E3 Ubiquitin-Protein Ligase Parkin and PTEN induced putative kinase (Pink1), both of them are associated with UPS. As inhibition of mTORC1 has been previoulsy shown to result in increased overall protein degradation *via* UPS and as FH mRNA and protein levels were shown to be affected by inhibition of UPS, our data stress a potential regulatory link between endogenous FH activity and the UPS.

## 1. Introduction

The ability to see is one of the most important faculties of the human body and visual impairment is reported to decrease the quality of life even more than chronic conditions like type II diabetes, hearing impairments or coronary disease [1]. In developed countries, age-related macular degeneration (AMD) has emerged to be one of the leading causes of legal blindness among the elderly [2–5]. Due to demographic changes and the strong association with age, AMD incidence is expected to strongly increase with a projected 288 million cases by 2040 [6]. In this light, AMD not only threatens a patient’s individual quality of life, but also poses a significant burden to health care systems worldwide [7,8]. AMD itself is a complex disease impacted by a wide range of causes, including genetic, life-style associated and environmental factors, which clinically manifests as a progressive atrophy of the central retina. Although the pathogenesis of AMD is still insufficiently understood, dysfunction of the Retinal Pigment Epithelium (RPE) plays a central role and is accompanied by degenerative processes in the retina, Bruch’s membrane (BrM) and the choriocapillaris (CC), ultimately leading to photoreceptor cells loss and consecutive vision impairment [9]. The RPE is a cellular interface between the retina and the BrM/CC that fulfills a plethora of tasks to maintain a physiological environment within the outer retina [10]. Accordingly, RPE cells are responsible for nutritional supply, waste disposal, recycling of shed outer segments and elimination of reactive oxygen species (ROS). To cope with these demands, RPE cells have to be highly metabolically active and even modest reductions in functionality may lead to cumulative, long-term damage to the outer retina [11]. Some of the most important risk factors for AMD, like age or smoking, increase ROS production that leads to excessive oxidative stress and in turn directly damages cellular structures and induce inflammatory processes [12]. In line with this, RPE cells from AMD patients show increased ROS production and vulnerability towards chronic oxidative stress [13]. To be able to recycle the large number photoreceptor shed outer segments accruing every day, the RPE heavily relies on phagocytosis in combination with central catabolic processes like autophagy and proteasomal degradation pathways [14,15]. Importantly, excessive oxidative stress has been linked to attenuated phagocytic potential of RPE cells and decreased autophagy was reported in RPE cells of AMD patients [13,16,17]. Those catabolic pathways are essential parts of cellular metabolism and the defects observed in AMD affect not only those, but metabolism as a whole. Profound dysregulation of central metabolic pathways, especially in those holding relevance for oxidative stress responses, has been observed in primary RPE cells obtained from AMD donor eyes [18].

One major regulator of cellular metabolism is the metabolic master-switch mammalian target of rapamycin (mTOR). In mammalian cells, mTOR is part of two distinct complexes: mTOR complex 1 (mTORC1) and mTOR complex 2 (mTORC2). mTORC1 works as a central integration hub that balances between anabolic processes and catabolic processes in respect to nutrient availability. In situations of nutrient availability, that allow for cellular growth, mTOR will block catabolic processes, like autophagy, and induce anabolic processes *via* direct phosphorylation of the main downstream effectors p70 ribosomal protein S6 kinase (S6K) and Eukaryotic translation initiation factor 4E-binding protein 1 (4E-BP1) [19]. Increased mTORC1 signaling is associated with cellular senescence and ageing [20], while mTOR inhibition is among the few interventions that ameliorates ageing effects and leads to life span extension in several model organisms [21]. In line with this, increasing the mTOR activity in a transgenic mouse model was sufficient to induce retinal degeneration [22]. Functional mTOR complexes (mTORC1 and mTORC2) are present in RPE cells [23] and increased mTOR activity was found in RPE cells of patients suffering from AMD [18]. Despite all evidences for a role of mTOR signaling in AMD, mechanistic insights on a molecular level are scarce but remain obligatorily for a rational development of novel and effective treatment strategies.

While environmental, nutritional and behavioral factors largely contribute to AMD pathogenesis, genetic predisposition also plays a prominent role in AMD development and progression. A large genome wide association study (GWAS) was able to detect 52 genetic risk variants in 34 loci that are associated with increased susceptibility for AMD development and progression [24]. Most of the highly associated variants map to genes related to the alternative complement pathway, an evolutionarily old part of the innate immune system involved in the neutralization of pathogens and dead or infected cells [25]. One of the most important complement-related AMD-associated SNPs (rs1061170) is located in the *CFH* gene, which codes for complement factor H (FH) and its truncated variant FHL-1 [26,27]. This polymorphism causes an amino acid exchange of Tyrosine at position 402 with a Histidine (Y402H; position 384 in the mature, secreted FH protein) [28]. Thus, the FH 402H variant is believed to contribute to AMD progression *via* uncontrolled complement system activation in the extracellular matrix [29]. Indeed, the FH 402H variant alters FH’s binding affinity towards polyanionic molecules, which in turn affects its ability to modulate complement activiation in the macula [30]. In addition, several *in vitro* studies suggest that FH holds additional functions in RPE cells, like regulation of cellular energy metabolism, which are affected by the AMD-associated variant [31] or by reduced intracellular FH levels and activity [32]. In particular, in our previous studies we assessed the impact of reduced levels of functional FH in RPE cells and found that reduced FH levels are associated with increased inflammatory signaling *via* the NF-κB pathway [33] and profound metabolic disturbances along with mitochondrial damage [31,32]. The exogenous supplementation of recombinant FH was not able to reverse those effects, pointing towards a non-canonical role of endogenous FH. Importantly, mTOR signaling plays a role in all of these processes: activation of the NF-κB pathway [34] and regulation of mitochondrial function [35] as well as cellular metabolism [36]. However, the mechanism by which intracellular FH modulates energy metabolism of RPE cells and whether mTOR is involved in this process is not known.

Therefore, this study investigates whether FH knockdown has an effect on mTOR activity and as a consequence, leads to metabolic damage. In parallel, we aimed to obtain insights into the mechanism of action of intracellular FH *via* analyzing the intracellular interactome of FH. Upon silencing of the *CFH* gene *via* RNAi to reduce FH protein levels in hTERT-RPE1 cells, we observed increased mTOR activity, demonstrated by increased mTOR phosphorylation (S2448) and increased phosphorylation of its main downstream effector protein, S6K (T389). Seahorse metabolic flux analysis revealed that mTOR inhibition *via* rapamycin is partially rescuing the defects in cellular respiration induced by FH knockdown, while glycolytic deficits remain largely unaffected. Mass-spectrometry-based analysis revealed that intracellular FH is predominantly interacting with factors associated with proteasomal protein degradation and cell cycle control. In line with previous results, this study proves the impact of reduced levels of intracellular FH on mTOR activity and point towards previously unknown intracellular functions of FH that will inform future studies to advance our mechanistic understanding of AMD pathophysiology.

## 2. Materials and Methods

### 2.1. Cell culture and experimental settings

The immortalized human retinal pigment epithelium (RPE) cell line hTERT-RPE1 was obtained from the American Type Culture Collection (ATCC). The cells were maintained in high-glucose Dulbecco’s modified Eagle’s medium (DMEM; Gibco, Waltham, MA, USA) supplemented with 10% fetal calf serum (FCS; Gibco, Waltham, MA, USA), penicillin (100 U/ml; Gibco, Waltham, MA, USA), streptomycin (100 µg/ml; Gibco, Waltham, MA, USA) in a humidified atmosphere containing 5% CO_2_.

Cells were seeded in 24-, 12- or 6-well plates or 10cm^2^ culture plates (Sarstedt, Nümbrecht, Germany) and allowed to attach overnight. Gene silencing was performed using Lullaby transfection reagent (OZ Biosciences, Marsaille, France) according to the manufacturer’s instructions using a equimolar mixture of three different double strand hairpin interference RNAs specific for *CFH* or a negative control as recommended by the provider (IDT technologies, Leuven, Belgium). Culture medium was substituted with fresh medium without antibiotics and siRNA mixture was added dropwise. After 24h incubation (37°C, 5% CO_2_), cells were switched to serum free medium (SFM) for 48 hours. In experiments including rapamycin treatment, cells were treated with DMSO (control) or rapamycin (Sigma-Aldrich; St. Louis, MO, USA) at the indicated concentrations. When indicated, cell culture medium was supplemeted with glucose (4,5g/l; AppliChem, Darmstadt, Germany), fructose (4,5g/l; Sigma-Aldrich; St. Louis, MO, USA) or amino acids (MEM amino acids solution; Gibco, Waltham, MA, USA).

### 2.2. RNA extraction, cDNA synthesis and quantitative RT-PCR

Total RNA was extracted using PureZOL reagent (Bio-Rad Laboratories, Des Plaines, IL, USA), according to the manufacturer’s instructions. Subsequent cDNA synthesis was done via reverse-transcription of 2µg of isolated RNA using M-MLV Reverse Transcriptase (Promega, Madison, WI, USA). The generated cDNA was used to analyze differences in gene expression by RT-qPCR employing iTaq Universal SYBR Green Supermix (Bio-Rad Laboratories, Des Plaines, IL, USA) along with gene specific forward and reverse primers (both 10 µM; purchased from IDT technologies, Leuven, Belgium) listed in Table 1. The used two-step PCR protocol consists of 40 cycles, each with 5 seconds at 95°C followed by 30 seconds at 57°C, run on a CFX96 Real-Time System (Bio-Rad Laboratories, Des Plaines, IL, USA). Relative mRNA expression of each gene of interest (GOI, Table 1) was quantified using 60S acidic ribosomal protein P0 (*RPLP0*) as the housekeeping control gene.

**Table 1:**
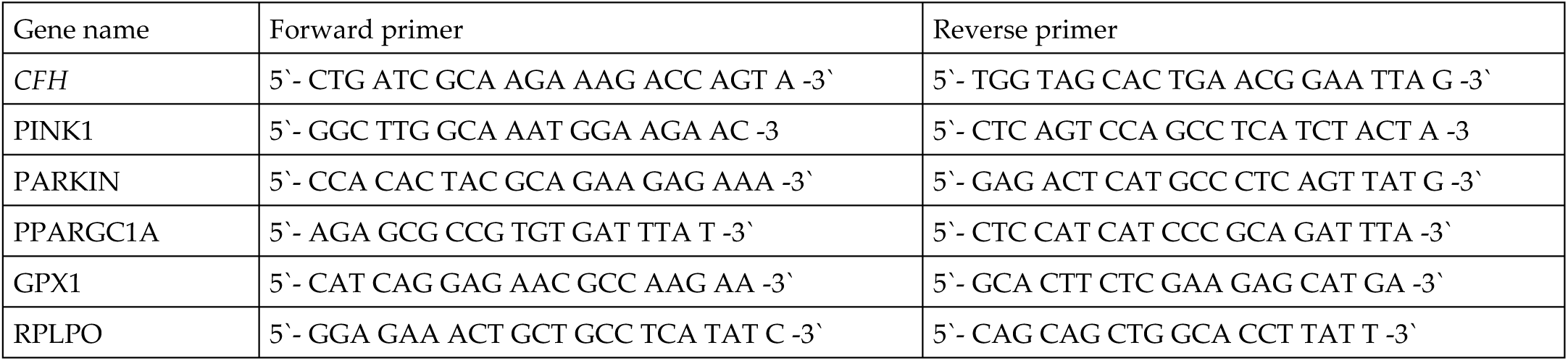
Primers list

### 2.3. Western Blotting

Protein expression was analyzed in both cell lysates and cell supernatants as previousely described [33]. Briefly, after debris removal, cell culture supernatants were precipitated with ice-cold acetone and cell lysates were prepared in Pierce IP Lysis Buffer, containing Halt Protease and Phosphatase Inhibitor (Thermo Fisher Scientific, Boston, MA, USA). Protein concentrations were determined with the Bradford quantification assay (Bio-Rad Laboratories, Des Plaines, IL, USA), using BSA (Sigma-Aldrich; St. Louis, MO, USA) as a standard. Equal protein amounts of cell lysates or equal volumes of cell supernatants were prepared in NuPAGE LDS Sample Buffer (Thermo Fisher Scientific, Boston, MA, USA), containing reducing agent (Thermo Fisher Scientific, Boston, MA, USA) and analyzed on Novex 8–16% Tris-Glycine gels (Invitrogen, Waltham, MA, USA). Subsequently, proteins were transferred onto PVDF membranes (Roche, Basel, Switzerland), and Western blot detection was carried out as previously described [32,33], using the primary antibodies listed in Table 2. Pictures were acquired with a FusionFX imaging system (Vilber Lourmat, Collégien, France), and the intensity density of individual bands was quantified using ImageJ (Version 1.53a).

**Table 2:** Antibody list

| Antibody | Supplier | Number |
| --- | --- | --- |
| Factor H (FH) | SantaCruz Biotechnology | sc-166608 |
| Phospho-mTOR (S2448) | Cell Signaling | #5336 |
| mTOR | Cell Signaling | #2983 |
| Phospho- S6K (T389) | Cell Signaling | #9234 |
| S6K | Cell Signaling | #9202 |
| β-actin | Cell Signaling | #3700 |

### 2.4. Immunoprecipitation and Mass-Spectrometry (MS)

Cells were grown on 10 cm^2^ dishes and after 48h of incubation in SFM, cell lysates were prepared using the IP lysis buffer (0.5% NP40 (Roche, Basel, Switzerland), 1x cOmplete protease inhibitor (Roche, Basel, Switzerland), 1% Phostphatase Inhibitor Cocktail 2 (Sigma-Aldrich; St. Louis, MO, USA), Phostphatase Inhibitor Cocktail 3 (Sigma-Aldrich; St. Louis, MO, USA) 1x TBS) and protein concentration was quantified using Bradford assays. Equal protein amounts were used for all cell lysates and volumes were adjusted using IP lysis buffer. Immunoprecipitation was performed using an anti-FH antibody (200µg/ml; santa-cruz, California, USA) or immunoglobulin G (200µg/ml; santa-cruz, California, USA) as a control. Lysates were incubated with the antibodies for 1h at 4°C on a roller before 80µl Protein G PLUS-Agarose beads (Santa Cruz Biotechnology, Dallas, TX, USA) were added and samples were incubated overnight at 4°C on a roller. Beads were pelleted (1000 g, 4°C, 5min), resuspended in IP washing buffer (0.1% NP40 (Roche, Basel, Switzerland), 1% Phostphatase Inhibitor Cocktail 2 (Sigma-Aldrich; St. Louis, MO, USA), Phostphatase Inhibitor Cocktail 3 (Sigma-Aldrich; St. Louis, MO, USA) 1x TBS) and transferred to 35µm receiver columns (Macherey-Nagel, Düren, Germany). After two washing steps using IP washing buffer, acidic elution with glycine buffer (200mM, pH=2) and neutralization with TRIS buffer (1M, pH=8.5) was performed. Proteins were precipitated using methanol/chloroform precipitation as previously described [37]. Elutaed proteins were processed and prepared for MS analysis as previously described [38]. MS analysis was performed on an Ultimate3000 RSLC system coupled to an Orbitrap Fusion Tribrid mass spectrometer (Thermo Fisher Scientific, Boston, MA, USA). Tryptic peptides were loaded onto a nano-trap column (300μm i.d. × 5mm precolumn, packed with Acclaim PepMap100 C18, 5μm, 100Å; Thermo Fisher Scientific, Boston, MA, USA) at a flow rate of 30 µl/min in 0.1% trifluoroacetic acid in HPLC grade water. After 3 minutes, peptides were eluted and separated on the analytical column (75μm i.d. × 25cm, Acclaim PepMap RSLC C18, 2μm, 100Å; Thermo Fisher Scientific, Boston, MA, USA) by a linear gradient from 2% to 30% of buffer B (80% acetonitrile and 0.08% formic acid in HPLC-grade water) in buffer A (2% acetonitrile and 0.1% formic acid in HPLC-grade water) at a flow rate of 300nl/minute over 150 minutes. Remaining peptides were eluted by a short gradient from 30% to 95% buffer B in 10 minutes. MS parameters were as follows: for full MS spectra, the scan range was 335–1,500 with a resolution of 120,000 at m/z=200. MS/MS acquisition was performed in top speed mode with 3 seconds cycle time. The maximum injection time was 50ms. The AGC target was set to 400,000, and the isolation window was 1 m/z. Positive Ions with charge states 2-7 were sequentially fragmented by higher energy collisional dissociation. The dynamic exclusion duration was set to 60 seconds and the lock mass option was activated and set to a background signal with a mass of 445.12002.

### 2.5. Mitochondrial respiration

Mitochondrial function was assessed in live cells using an XFp Extracellular Flux Analyzer (Agilent Technologies, Santa Clara, CA, USA). After 24 hours silencing (siNeg vs siCFH) in 6-well-plates, hTERT-RPE1 cells (3 × 10^4^ cells/well) were seeded in at least duplicates in XFpSeahorse microplates and allowed to adhere overnight. Following medium change, cells were grown for further 48 hours in medium containing DMSO (as control) or Rapamycin resolved in DMSO. DMSO concentrations did not excede 0.1% of the medium. Measurements of oxygen consumption rate (OCR) were performed in freshly prepared assay medium, pH 7.4 (Seahorse XF DMEM Medium), according to the manufacturer’s protocol. OCR was measured before and after the serial addition of inhibitors: 1) 10 μM Oligomycin, 2) 10 μM carbonyl cyanide p-trifluoromethoxyphenylhydrazone (CCCP) and 3) 1 μM Antimycin A and 1 μM Rotenone. to assess several parameters of mitochondrial function (Figure 2a). Following each injection, 3 measurements for a total period of 15 minutes were recorded. The data were analyzed using Wave 2.6 Software and OCR parameters (basal respiration, maximal respiratory capacity, respiratory reserve and ATP-linked respiration) were calculated. All reagents were purchased from Sigma-Aldrich; St. Louis, MO, USA. At least 2 technical replicates per condition were used and the experiemental values from 4-5 independent experiments were normalized to protein content/well *via* BCA assay (Pierce Biotechnology, Waltham, MA, USA).

### 2.6. Glycolysis

Glycolysis function was assessed in live cells using an XFp Extracellular Flux Analyzer (Agilent Technologies, California, USA). After 24 hours silencing (siNeg vs siCFH) in 6-well-plates, hTERT-RPE1 cells (3 × 104 cells/well) were seeded in at least duplicates in XFpSeahorse microplates and allowed to adhere overnight. Following medium change, cells were grown for further 48 hours in medium con taining DMSO or Rapamycin. Measurements of extra-cellular acidification rate (ECAR) were performed in freshly prepared assay medium, pH 7.4 (Seahorse XF DMEM Medium), according to the manufacturer’s protocol. ECAR was measured before and after the serial addition of 1) 10 mM glucose, 2) 10 µM oligomycin and 50 mM 2-deoxy-glucose (2-DG). Following each injection, 3 measurements for a total period of 15 minutes were recorded. The data were analyzed using Wave 2.6 Software and ECAR parameters (basal glycolyis, glycolytic capacity and glycolytic reserve) were calculated. All reagents were purchased from Sigma-Aldrich; St. Louis, MO, USA. At least 2 technical replicates per condition were used and the experiemental values from 4-5 independent experiments were normalized to protein content/well *via* BCA assay (Pierce Biotechnology, Waltham, MA, USA).

### 2.7. Bioinformatic anaylsis of MS-data

Analysis of MS data was performed using the MaxQuant [39] software (version 1.6.17.0). Trypsin was selected as the digesting enzyme with maximal 2 missed cleavages. Cysteine car-bamidomethylation was set for fixed modifications and oxidation of methionine and N-terminal acetylation were specified as variable modifications.. The first search peptide tolerance was set to 20, the main search peptide tolerance to 5ppm. For peptide and protein identification the Human subset of the SwissProt database (Release 2021_04) was used, and contaminants were detected using the MaxQuant contaminant search. A minimum peptide number of 1 and a minimum length of 6 amino acids was tolerated. Unique and razor peptides were used for quantification. The match between run option was enabled with a match time window of 0.05 min and an alignment time window of 20 min. The statistical analysis including ratio and two sample t-test was done using Perseus [40]. Identification of FH potential interactors was performed using a one sided permutation based t-test with 250 randomizations, an FDR < 0.05 and a S0 of 0.1.

### 2.8. Statistical analyses

The data are presented as mean with the standard error of the mean (SEM) and were generated and tested for their significance with GraphPad Prism 8 software (Version 8.4.3). All data sets were tested for normal distribution, assessed with the Shapiro-Wilk normality test. Depending on the normal distribution and the parameters to be compared, the following tests were performed. A paired Student’s t-test was used to compare siNeg vs siCFH conditions, siNeg vs siNeg + Rapa conditions, siCFH vs siCFH + Rapa conditions. Values were considered significant with a p-value < 0.05.

## 3. Results

### 3.1. FH knockdown leads to increased mTOR activity

To assess the impact of decreased levels of FH on mTOR activity, *CFH* was silenced *via* RNAi and silencing efficiency was verified at both the RNA level, using qPCR (Figure S1a), and at the protein level, *via* western blot (Figure S1b). Across all experiments, RNA levels were reduced by 80% and FH protein levels were reliably reduced by more than 80% in siCFH cells relative to control cells (Figure S1a-b). To investigate the activation levels of the mTOR pathway, we monitored the total mTOR protein levels and levels of the activated phosphorylated form of mTOR at serine 2448 (Figure 1a), a well-established marker for i*n vivo* mTOR activity. We found a significant increase in mTOR activation levels in *CFH* silenced cells (siCFH) compared to control cells (siNeg), as shown by the increased ratio of phosphorylated/total mTOR (**Error! Reference source not found**.a). In accordance with mTOR phosphorylation, we observed significantly increase S6K activation levels, as shown by the increased ratio of phosphorylated/total S6K (**Error! Reference source not found**.b). As mTOR serves as a molecular integration hub for numerous metabolically relevant stimuli, the impact of several nutritional conditions on mTOR activity in siNeg and siCFH cells were analyzed. To mimic nutrient scarcity and physiologically decreased mTOR activity, serum-free media lacking glucose (-GLC) or Glutamine (-GLN) were used. To mimic nutrient availability and to physiologically upregulate mTOR activity, serum-free medium containing elevated levels of amino acids (hAA) was used. Additionally, glucose was substituted with fructose (+FRC) to resemble the high fructose content of typical western-type diets that are typically associated with increased risk for AMD. Generally, the siCFH cells showed higher levels of active mTOR in most of the conditions (ctrl, -GLC, +FRC) and minimal changes when the levels of essential amino acid as glutamine were reduced (-GLN) or increased (hAA) (**Error! Reference source not found**.c). Then, siCFH cells continuously showed a trend towards higher S6K phosphorylation compared to siNeg cells, with differences reaching significance when incubated with SFM and –GLC (Figure 1d). Overall, the different nutritional conditions – except - GLC - had no major impact and mTOR activity.

**Figure 1:**
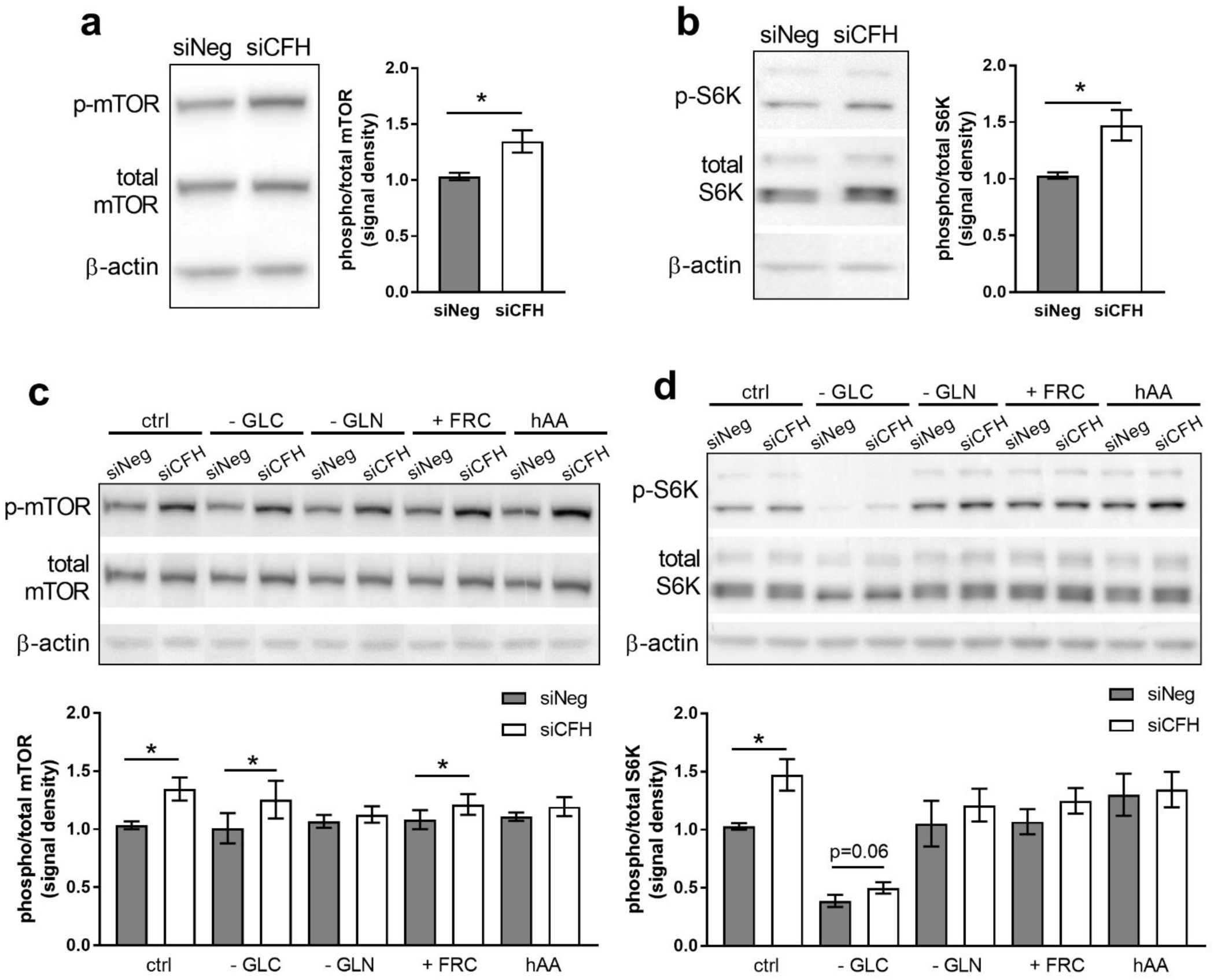
FH knockdown leads to mTOR pathway activation in RPE cells. hTERT-RPE1 cells were silenced for 24 hours with negative control (siNeg) or CFH specific (siCFH) siRNA and then incubated for 48 hours in the indicated conditions. Cell pellets were collected for protein extraction. a,c) Western blot analyses of phosphorylated and total levels of mTOR. Total β-actin was used as loading control. Quantification of at least 4 independent experiments is shown, where bars indicate the signal density ratio between levels of phosphorylated and total mTOR. b,d) Western blot analyses of phosphorylated and total levels of S6K. Total β-actin was used as loading control. Quantification of at least 4 independent experiments is shown, where bars indicate the signal density ratio between levels of phosphorylated and total S6K. SEM is shown. Significance was assessed by Student’s t-test. *p < 0.05. GLC, glucose. GLN, glutamine. FRC, fructose. hAA, high aminoacids.

### 3.2. mTOR inhibition via Rapamycin partially reverses FH knockdown mediated defects in cellular respiration

As previously reported, hTERT-RPE1 cells silenced for *CFH* (siCFH) show drastically reduced levels of mitochondrial respiration and glycolysis, analyzed *via* the oxygen consumption rates (OCR) and extracellular acidification rates (ECAR), respectively, in Seahorse metabolic flux analyses [32]. To assess whether FH knockdown mediates its effects on cell metabolism *via* overactivation of the mTOR pathway, we treated siNeg and siCFH RPE cells with rapamycin, a well-known inhibitor of mTORC1 activity. The inhibitory capacity of rapamycin in hTERT-RPE1 cells was confirmed by analyses of the phosphorylation levels of S6K, a main target of mTORC1-mediated phosphorylation (Figure S2a). Western blot analyses showed that concentrations ranging from 0.1 to 200nM were sufficient to completely abolish S6K phosphorylation (Figure S2a) without significantly impacting on cell viability (Figure S2b). Based on the good tolerability and strong inhibition of mTOR activity, a final rapamycin concentration of 200nM was chosen for further experiments, while controls were treated with vehicle (DMSO) only. In line with previous experiments, untreated siCFH cells showed significantly reduced OCR rates compared to siNeg cells (**Error! Reference source not found**.a), with basal respiration reduced by 37 % (Figure 2b), maximal respiration by 50 % (Figure 2c), reserve respiratory capacity by 56 % (Figure 2d) and ATP-linked respiration by 38 % (Figure 2e). Rapamycin treatment in siCFH RPE cells resulted in a small but significant increase in the OCR parameters compared to untreated siCFH RPE cells (Figure 2a): 22 % for basal respiration (Figure 2b), 35 % for maximal respiration (Figure 2c), 32 % for reserve respiratory capacity (Figure 2d) and 25 % for ATP-linked respiration (Figure 2e). No significant differences were observed between untreated siNeg cells and Rapamycin treated siNeg cells (**Error! Reference source not found**.b-e). Analysis of the extracellular acidification rate (ECAR) showed significant reductions in siCFH cells relative to siNeg cells but rapamycin treatment did not have any effects on the glycolytic parameters (Figure S3).

**Figure 2:**
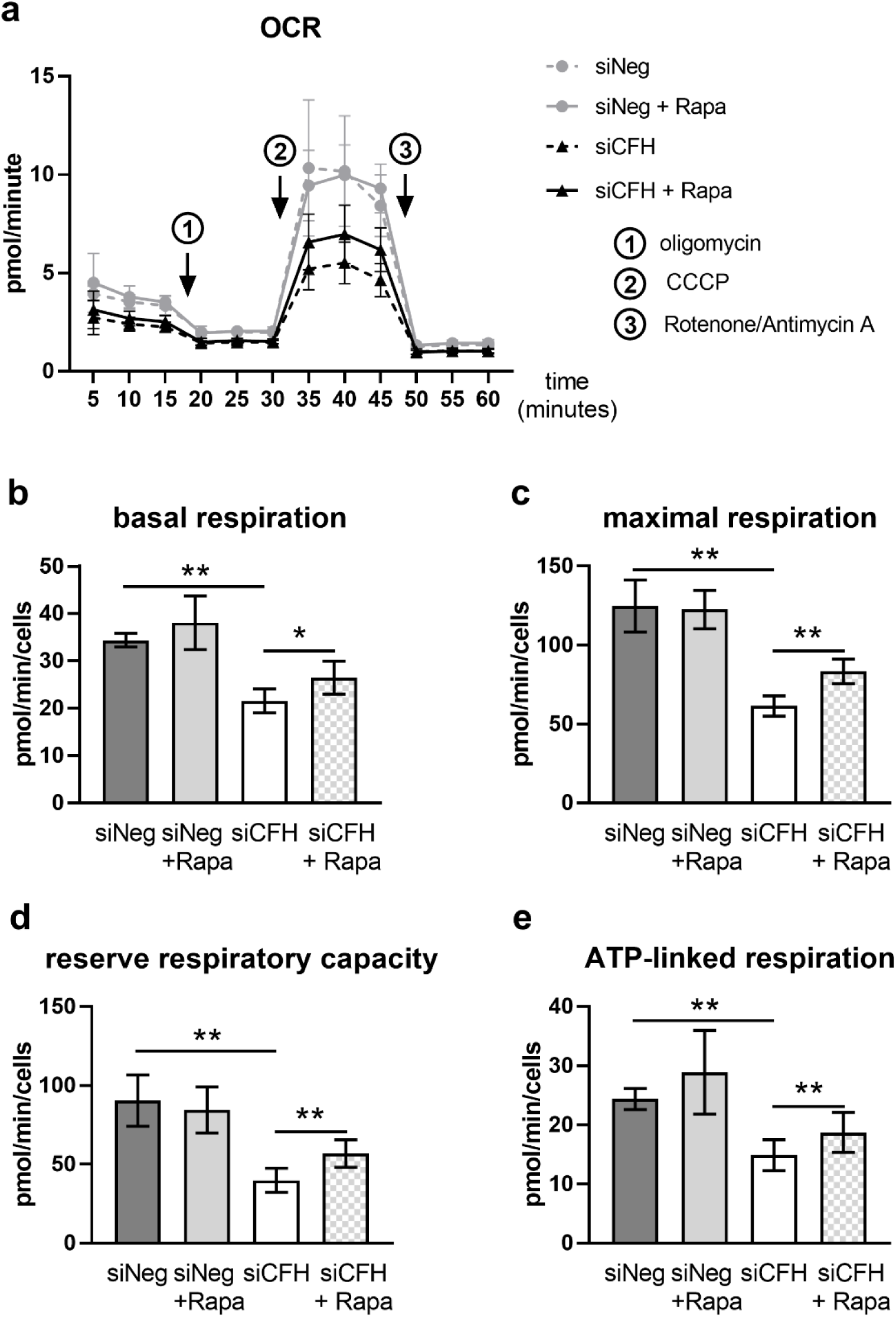
Rapamycin partially rescues FH knockdown-mediated mitochondrial respiration impairment in RPE cells. (a) hTERT-RPE1 cells were silenced for 24 hours with negative control (siNeg) or CFH specific (siCFH) siRNA, seeded on seahorse plates and incubated in SFM containing 200nM rapamycin or DMSO. Curves represent the oxygen consumption rates (OCR) measured after 48 hours. SEM shown, n = 4–5. Arrows indicate injection of oligomycin (1), CCCP (2) and antimycin and rotenone (3). (b–e) Parameters of mitochondrial respiration were calculated based on the data shown in (a). (b) Basal respiration, (c) maximal respiration, (d) reserve respiratory capacity and (e) ATP-linked respiration. Significance was assessed by Student’s t-test. *p < 0.05, **p < 0.01.

### 3.3. mTOR inhibition via Rapamycin reverses FH knockdown mediated effects on gene expression

In our previous study, we show that in siCFH RPE cells gene expression of several factors involved in mitochondria stability and antioxidant response were upregulated [32]. Here, we evaluated whether those genes were downstream targets of m TOR activation. First, we evaluated the gene expression levels of genes regulating mitophagy: E3 Ubiquitin-Protein Ligase Parkin (PARKIN, Figure 3a) and PTEN induced putative kinase 1 (PINK1, Figure 3b). We confirmed the upregulation of both genes in siCFH compared to siNeg cells, although here more pronounced for PARKIN than as previousely described [32]. Rapamycin treatment was able to revert the upregulation of both genes, when comparing siCFH cells versus siCFH+Rapa conditions (Figure 3a-b). Then, we monitored the gene expression levels of Peroxisome Proliferator-Activated Receptor Gamma Coactivator 1-Alpha (PGC1α, Figure 3c), a transcription factor promoting mitochondria biogenesis, which is often activated in case of mitochondrial damage. We confirm the upregulation of PGC1α in siCFH compared to siNeg cells and that rapamycin treatment leads to a significant reduction of the gene expression levels of PGC1α (siCFH vs siCFH +Rapa, Figure 3c). Lastly, we monitored the changes in Glutathione Peroxidase 1 (GPX1, Figure 3d), an antioxidant enzyme that is induced by PGC1α. The RNA levels of GPX1 were significantly increased in siCFH cells compared to siNeg cells and the effects were reverted by rapamycin treatment.

**Figure 3:**
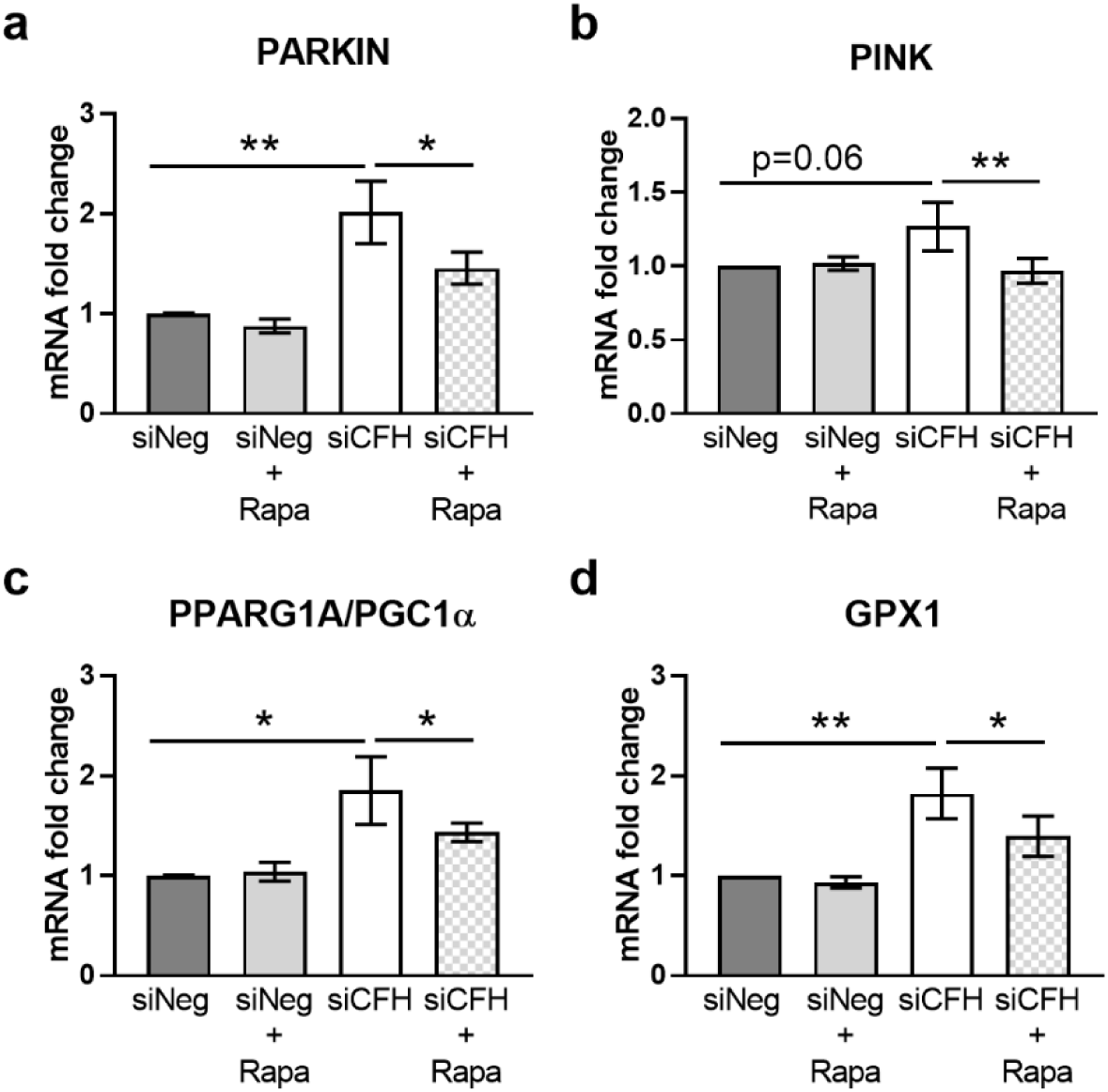
Rapamycin reverts FH knockdown-mediated effects on gene expression. hTERT-RPE1 cells were silenced for 24 hours with negative control (siNeg) or CFH specific (siCFH) siRNA and then incubated in SFM containing 200nM rapamycin or DMSO for 48 hours. Cell pellets were collected for RNA extraction, cDNA synthesis and qRT-PCR analyses. a) Evaluation of gene expression levels of E3 Ubiquitin-Protein Ligase Parkin (PARKIN). b) Evaluation of gene expression levels of PTEN induced putative kinase 1 (PINK1). c) Evaluation of gene expression levels of Peroxisome Proliferator-Activated Receptor Gamma Coactivator 1-Alpha (PGC1α). d) Evaluation of gene expression levels of Glutathione Peroxidase 1 (GPX1). Data are normalized to housekeeping gene RPLP0 using ΔΔCt method. SEM is shown, n=8. Significance was assessed by Student’s t-test. *p < 0.05, **p < 0.01.

### 3.4. Intracellular FH physically interacts with components of the proteasome and factors involved in cell cycle regulation

Mass-spectrometry-based analysis (Figure 4A) of the intracellular FH-interactome yielded a set of 101 identified proteins with significant enrichment in the FH pulldown samples in comparison to the IgG control samples (Table S1). A gene-ontology enrichment analysis using the KEGG 2021 database *via* Enrichr [41] identified the proteasomal pathway and cell cycle as the most represented pathways (Figure 4B). Protein-protein interaction network analysis (PPI) using metascape [42] revealed two distinct clusters of interacting proteins. FH was found to interact with several components of the 20S/26S proteasomes (PSMD13, PSME3, PSMB3, PSMB6, PSMA3) and with factors associated with the Rb1/E2F pathway (E2F3, E2F4, RB1, TFDP1, TFDP2) (Figure 4C).

**Figure 4:**
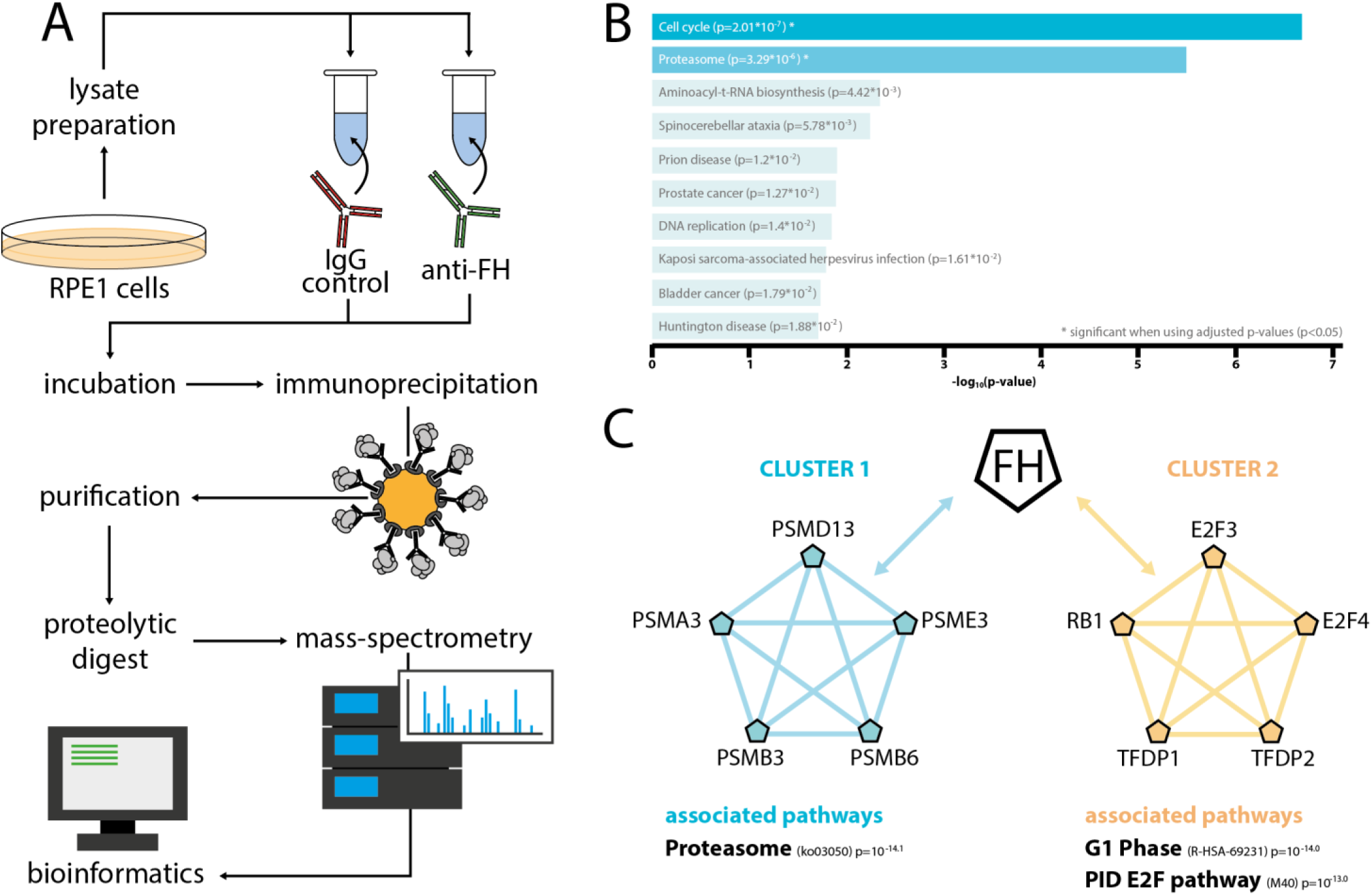
intracellular FH interacts with facotrs associated with the proteasomal pathway and the RB1/E2F pathway. hTERT-RPE1 lysates were incubated with either unspecific immunoglobulin G (control) or anti-FH antibody and immunoprecipitation was perfromed using protein-G labeled agarose beads. Proteins were purified, digested with trypsin before mass-sprectrometric analysis. a) Experimental workflow. b) Gene-ontology analysis using Enrichr [41] identified the proteasomal pathway and cell cycle as the most represented pathways. c) Protein-protein interaction network analysis via Metascape [42] defined two clusters of FH interacting factors. Cluster 1 included members of the 20S/26S proteasome (PSMD13, PSME3, PSMB6, PSMB3, PSMA3), cluster 2 included factors involved in RB1/E2F signaling and cell-cylce control (E2F3, E2F4, TFDP2, TFDP1, RB1).

## 4. Discussion

As a disease involving several cell types in different tissues, elucidation of AMD-pathology has been and continues to be a significant challenge for the scientific community. The complex interplay of numerous factors including genetic predisposition, environmental and life-style-associated factors along with the strong association with aging, make it difficult to draw definite conclusions and develop effective treatment strategies. AMD-related research is largely limited by the absence of well-suited models and close consideration needs to be paid when choosing the models to be used for a specific scientific question. While mouse models allow for certain AMD-related analyses, the fact that only humans and higher-order primates develop AMD and that the typical macular structure is absent in any lower-order model organism, is limiting the use of animal models. On the other hand, simpler models, like for example cell-cultures of immortalized AMD-relevant cell lines, are not able to resemble the complex interplay between different cell types and different tissues in the eye. Nevertheless, they may be well suited to analyze fundamental cellular processes and to identify potential targets for future studies in more complex models. In previous studies we and others could show, that hTERT-RPE1 cells feature physiological as well as pathological interrelationships also observed in more complex AMD-model systems. For example, the mitochondrial damage and impairment in cellular metabolism observed upon *CFH* silencing [32], closely resembled observations made in patient-derived iPSC-RPE cells carrying the AMD-associated FH risk variant (rs1061170) [31].

In this study, we used *CFH* silenced hTERT-RPE1 cells to analyze the impact of FH dysregulation on mTOR, a well-conserved pathway across species, tissues and cell types. *CFH* knockdown resulted in increased mTOR phosphorylation at Serine 2448 (Figure 1a,c), a marker for *in vivo* mTORC1 activity [43]. In line with increased mTOR activity, we observed increased phosphorylation of S6K at threonine 389 (Figure 1b,d), which is a direct target of mTORC1 kinase activity and a major effector of the mTOR pathway [44]. Significant differences between siCFH and siNeg cells in p-mTOR(S2448) and p-S6K(T389) were observed for most tested conditions. However, the observed differences have to be considered relatively small. Given the fact that mTOR integrates a huge number of different molecular stimuli [45] along with the fact that FH is not reported to be a direct mTOR interacting protein, it is expected that the impact FH dysregulation on overall mTOR activity may be limited. Nevertheless, the central role of mTOR in cellular metabolism and cellular homeostasis, render even subtle changes in mTOR activity physiologically relevant, especially in diseases that develop over a long period of time. In line with this, RPE cells from AMD-patients showed elevated levels of p-mTOR(S2448) and p-S6K(T389) compared to non-AMD RPE, with differences being in a similar range as observed in our study [18].

Furthermore, mTOR activity is closely linked to aging and age-related diseases [46]. AMD-related changes are also observed in eyes from healthy donors, but appear at later age and show slower rates of progression [10]. For instance, small drusen (<63µm in diameter, now commonly called drupelets) can be found in the non-AMD-related aging retina [47]. Similarily, slight, but constantly elevated mTOR activity together with an elevation of pro-inflammatory or stress related factors [31,33,48] may accelerate age-associated processes that push the development of AMD. Previous studies have shown that FH dysregulation in hTERT-RPE1 cells leads to increased vulnerability towards oxidative stress [32]. Furthermore, increased oxidative stress alone is sufficient to induce RPE dedifferentiation in wild-type mice *via* stimulation of the Akt/mTOR pathway and these detrimental effects were blunted by mTOR inhibition *via* rapamycin [49]. Based on the rapamycin’s proven ability to prevent or even reverse detrimental effects caused by elevated mTOR activity, we tested if it can reverse the metabolic deficits elicited by FH dysregulation. In accordance with our previous study [32], we employed metabolic flux analyses to monitor the effect of rapamycin treatment on oxygen consumption rates (OCR) (Figure 2) and extracellular acidification rates (ECAR) (Figure S3). As expected, cells treated with *CFH* specific siRNA (siCFH) showed drastically reduced OCRs (Figure 2) and ECARs (Figure S3) compared to siNeg cells. Rapamycin treatment lead to significant increases in OCR in siCFH but not siNeg cells. However, the defects were not completely rescued. This may be explained by the experimental conditions and it is possible that prolonged incubation intervals or dosage modulation would further improve the experimental outcome. Another reasonable interpretation is that hyperactive mTOR is only one of several pathological effects induced by FH dysregulation and that a complete rescue demands combinatorial interventions. In agreement with our findings, RPE-specific mTOR hyperactivation in a transgenic mouse model leads to retinal degeneration and rapamycin treatment only partially rescued the phenotype [22]. A phase II clinical trial assessing the impact of intravitreal rapamycin on disease progression in patients with geographic atrophy (GA) did not show any relevant differences between the treatment and control group [50]. However, the study is limited by the low number of included subjects and the advanced disease stage at treatment initiation. Furthermore, based on the multifactorial nature of AMD, sub-stratification based on clinical parameters and/or genetic risk status may be needed [51] as it is likely that different AMD-subgroups with differing molecular disturbances exist and that a one-size-fits-all approach is prone to yield suboptimal results. In concrete terms, this could mean, that stratification of patiens along FH dysregulation is the correct approach for rapamycin-mediated trials.

The fact that the exogenous supplementation of recombinant FH was not able to reverse the defects in cellular metabolism elicited by FH knockdown in RPE cells [32,33] points towards a non-canonical, intracellular, role of endogenous FH. Interestingly, the non-canonical intracellular role of FH has been recently described in other cell types, showing a similar phenotypes as RPE cells. For example, knockdown of FH impact several features of clear cell renal cell carcinoma cells *in vitro* in a complement cascade independent manner and transcriptomic analysis shows defects in cell cycle regulation [52]. To gain mechanistic insights, we performed a mass-spectrometry based analysis of the intracellular FH interactome. None of the known FH ligands, i.e. C3, FI, were found. This is not surprising, given that the intracellular microenvironment largely differs from the extracellular space in its specific biophysical conditions and molecular composition. Thus, FH is unlikely to bind any of these well-known extracellular interactors within the cell itself. With the differing sets of interacting proteins within and outside the cell, it is apparent that the intracellular function may be largely different from the known extracellular role of FH. Our compelling data notwithstding, it can not be entirely ruled out there may be a potential contribution of miRNA disruption on our observations. Two NF-κB regulated miRNAs, *i.e*. miRNA-125b and miRNA-146a, target *CFH* mRNA by binding the miRNA regulatory control region in the 3’-untranslated region (3’UTR) of the *CFH* mRNA [53]. It is concievable that by scilencing the *CFH* gene the miRNAs, that would otherwise bind the CFH mRNA 3’UTR, would instead bind to other targets. Interestingly, alternative binding partners of miRNA-125b include tumor necrosis factor alpha-induced protein 3 (TNFAIP3), which itself regulates the NF-κB signaling pathway [54], thus creating an NF-κB activation feedback loop. Similarly, miRNA-146a regulates not only inflammatory pathways, but has also been found to regulate metabolic pathways including mTOR in adipose tissue macrophages [55]. Although effects on the mRNA level cannot be ruled out at this point, it is to be considered unlikely as miRNA-146a knockdown mice showed increased mTOR singaling, while CFH knockdown should lead to stoichiometric increases in miRNA-146a availability and thus should decrease mTOR activity. In contrast, we observed mTOR upregulation upon CFH knockdown. Taken together, there remains no doubt that disturbance of the *CFH* gene in RPE cells adversley affects their health and abaility to support their adjacent neurosensory retina and demonstrates the need to better understand the retina as a multicellular tissue rather than isolated cell types.

Unexpectedly, the most prominent interactor patterns were made up by factors involved in the ubiquitin-proteasomal pathway (UPS) and central effectors of RB1/E2F signaling pathways (Figure 4). Besides autophagy, the UPS is the major intracellular system responsible for protein and organelle degradation [56]. Both processes are inherently interconnected and impairments in one pathway can directly alter the activity of the other [57], providing a potential link between mTOR, a major regulator of autophagy, and the UPS. Inhibition of autophagy in ARPE19 cells results in impaired UPS [58] and inhibition of mTORC1 leads to increased overall protein degradation *via* the UPS in HEK293 cells [59]. Intriguingly, ARPE19 cells showed decreased *CFH* mRNA and protein levels upon chemical inhibition of the UPS [60], providing a potential regulatory link between FH and the UPS. In addition to Ebeling et al., who speculated that FH dysregulation may indirectly impact on the proteasomal activity in RPE cells *via* the intracellular complement system [31], our data add another layer of complexity as they imply, that FH, endogenously produced by RPE cells, directly interacts with essential components of the UPS in a complement-independent manner. While a link between autophagy, the UPS and FH is evident, the exact mechanism of action of FH remains unclear and needs to be addressed in future studies.

RB1 is a well known tumor suppressor involved in CDK4/CDK6-dependent cell-cycle regulation. CDK4 and CDK6 phosphorylate RB1, cause the inactivation and de-repression of the RB1-associated E2F transcription factors. The de-repressed E2F transcription factors then induce numerous genes that drive cell-cycle progression [61,62]. Although past studies largely focussed on RB1’s role in cell-cycle regulation, more diverse, non-canonical roles, like in metabolism, have been reported [63]. Both, CRISPR-mediated knockout of RB1 as well as RB1 knockdown *via* RNAi in hTERT-RPE1 cells resulted in drastically decreased OCR, decreased epxression of mitochondrial proteins and mitochondrial activity [64]. Future experiments may therefore assess a direct potential interplay between FH and RB1 to provide a mechanistic explanation for the OCR decrease observed in FH knockdown RPE cells. Besides the metabolic involvement of RB1, its impact on cell-cycle regulation may be of relevance for AMD. If FH impacts RB1 activity, FH dysregulation may lead to cell-cycle dysregulation and contribute to RPE senescence, which has been associated with AMD pathogenesis [65].

Taken together, our findings on the *CFH* silencing RPE model allowed us to highlight the importance of FH on cellular homeostasis, including cellular metabolism and inflammatory response. Based on our data, we suggest the mTOR pathway as partially responsible for the mitochondrial damage caused by FH loss in RPE cells. Moreover, for the first time, we provide a panel of intracellular binding partners of FH, that may provide a basis for further research focused on elucidating the exact mechanism of action of FH inside the RPE cells and we provide the proteosomal pathway and RB1/E2F signaling as main candidates.

## Supplementary Materials

Figure S1: Efficiency of CFH silencing in RPE cells, Figure S2: effects of Rapamycin concentration on mTOR pathway and viability, Figure S3: Effects of rapamycin on the glycolysis of RPE cells, Table S1: Significantly enriched proteins in FH samples.

## Author Contributions

Conceptualization: A.A. and M.Ue; methodology: D.A.M., F.P., A.A. M.A.J.; software: D.A.M., F.P., A.A. M.A.J.; validation: D.A.M., F.P., A.A. M.A.J.; formal analysis: D.A.M., F.P., A.A. M.A.J.; investigation: D.A.M., F.P., A.A. M.A.J., M.Ue.; resources: A.A.; M.Ue.; S.J.C.; M.D..; data curation: D.A.M., F.P., A.A. M.A.J., E.K.; writing—original draft preparation: D.A.M., A.A.; writing—review and editing: D.A.M., F.P., A.A. M.A.J., E.K.; S.J.C.; M.D., M.Ue.; visualization: D.A.M., A.A., M.A.J.; supervision: A.A., S.J.C., M.D.; M.Ue.; project administration: E.K.; funding acquisition: A.A., M.Ue, S.J.C.; M.D.. All authors have read and agreed to the published version of the manuscript.

## Data Availability Statement

All raw data was uploaded to the ProteomeXchange database and will be made publically available upon publication of the manuscript.

