## supplementary material for "mTOR inhibition *via* Rapamycin treatment partially reverts the deficit in energy metabolism caused by FH loss in RPE cells"

*antioxidants*

*Research Article*

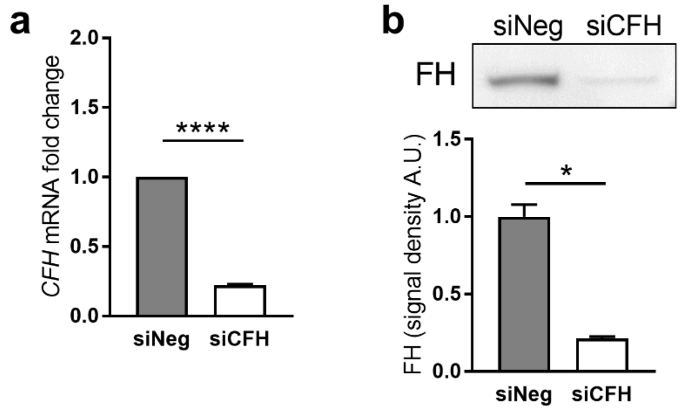


*Figure S1: Efficiency of CFH silencing in RPE cells*. hTERT-RPE1 cells were silenced for 24 hours with negative control (siNeg) or CFH specific (siCFH) siRNA and then incubated for 48 hours. Cell pellets and cell culture supernatants were collected for RNAand protein extraction. a) Evaluation of *CFH* expression by qRT-PCR analyses. Data are normalized to the housekeeping gene PRLP0 using Δ ΔCt methods. SEM is shown, n=4. b) Western blot analyses of FH protein levels in cell culture supernatants of hTERT-RPE1 cells. Quantification of signal density of 3 independent experiments is shown Significance was assessed by Student's t-test.

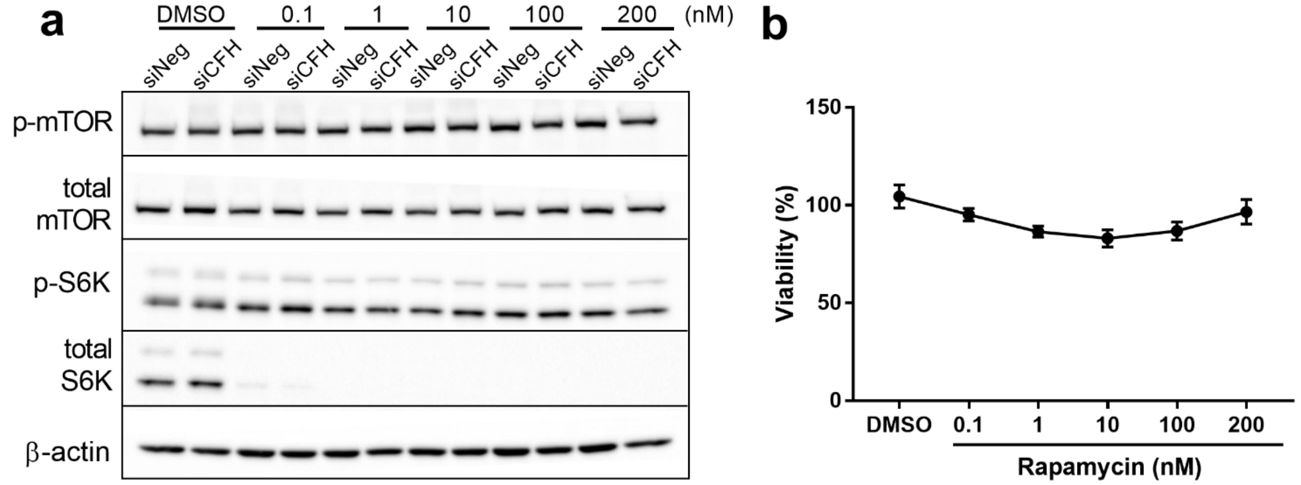


*Figure S2: effects of Rapamycin concentration on mTOR pathway and viability*. hTERT-RPE1 cells were silenced for 24 hours with negative control (siNeg) or CFH specific (siCFH) siRNA and then incubated for 48 hours with increasing concentrations of rapamycin as indicated. a) Cell pellets were collected for protein extraction. Western blot analyses of phosphorylated and total levels of mTOR; phosphorylated and total levels of S6K. Total β-actin was used as loading control. b) Viability assessed *via* MTT assay. N=3. SEM is shown.

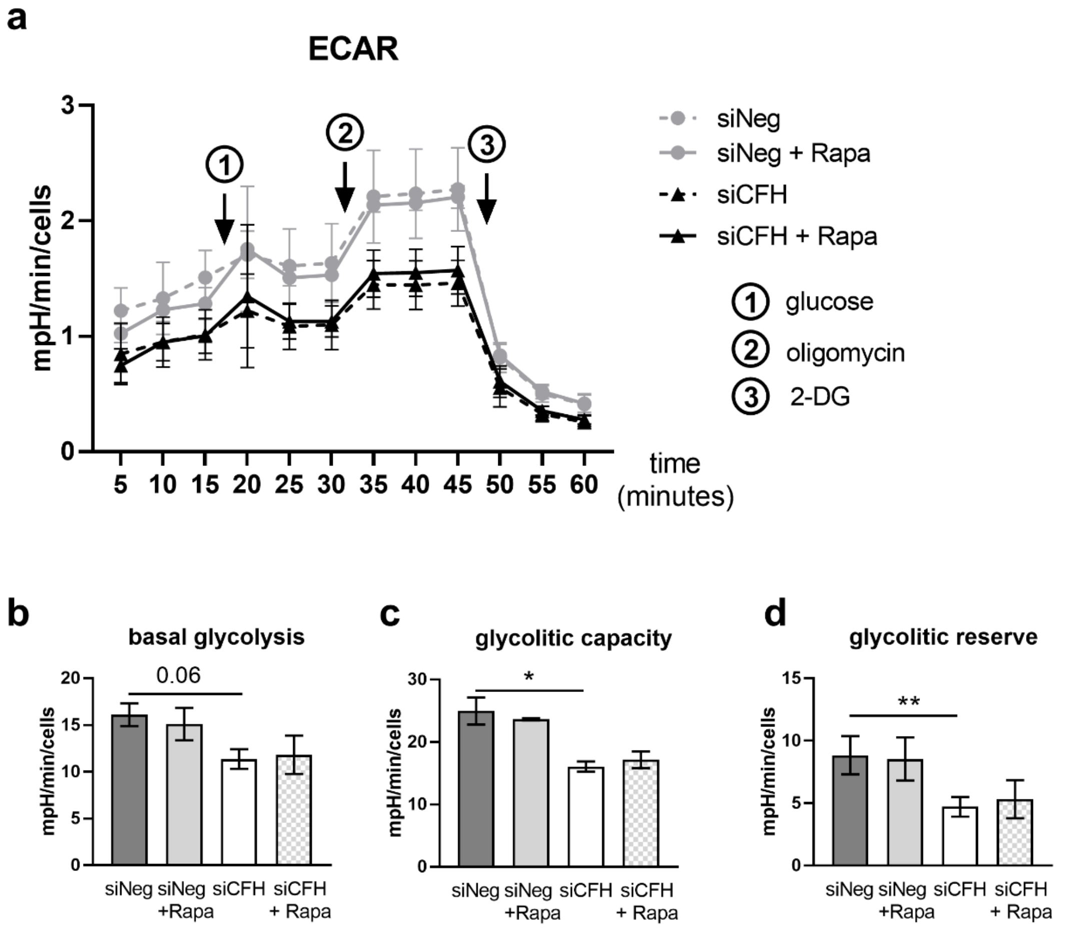


*Figure S3: Effects of rapamycin on the glycolysis of RPE cells*. (a) hTERT-RPE1 cells were silenced for 24 hours with negative control (siNeg) or CFH specific (siCFH) siRNA, seeded on seahorse plates and incubated in SFM containing 200nM rapamycin or DMSO. Curves represent the extracellular acidification rates (ECAR) measured after 48 hours. SEM shown, n = 5. Arrows indicate injection of glucose (1), oligomycin (2), 2-DG (3). (b–d) Parameters of glycolysis were calculated based on the data shown in (a). (b) Basal glycolysis, (c) glycolytic capacity, (d) glycolytic reserve. Significance was assessed by Student's t-test. *$p < 0.05$, **$p < 0.01$.

Table S1: Significantly enriched proteins in FH samples. The whole dataset including LFQ intensities and additional parameters can be retrieved from ProteomeXchange.

| Gene names | Majority protein IDs | p-Value | Mean Difference CFH vs Ctrl |
|---|---|---|---|
| **LEPRE1** | Q32P28 | 1.6235E-14 | 23.7274313 |
| **CCDC136** | Q96JN2 | 3.33475E-14 | 26.80236149 |
| **CREB1;ATF1** | P16220;P18846 | 4.99625E-14 | 24.20196581 |
| **BRAP** | Q7Z569 | 6.50194E-14 | 24.82372427 |
| **RABGGTB** | P53611 | 1.05303E-13 | 22.96165276 |
| **SPTB** | P11277 | 1.07642E-13 | 24.1661644 |
| **FAM184B** | Q9ULE4 | 1.10747E-13 | 29.97203302 |
| **E2F4** | Q16254 | 1.12392E-13 | 23.22352886 |
| **UCN3** | Q969E3 | 1.34454E-13 | 23.88911533 |
| **RAD17** | O75943 | 4.11222E-13 | 25.5679965 |
| **NCKAP5L** | Q9HCH0 | 9.4782E-13 | 21.06704617 |
| **ITCH** | Q96J02 | 1.9754E-12 | 23.4377985 |
| **ATAD3A** | Q9NVI7 | 4.30175E-12 | 21.26448727 |
| **LUC7L3** | O95232 | 5.21921E-12 | 21.21871471 |
| **KIAA1107** | Q9UPP5 | 7.15949E-12 | 20.27953434 |
| **GNB4** | Q9HAV0 | 8.33428E-12 | 20.82388973 |
| **CACTIN** | Q8WUQ7 | 8.67736E-12 | 21.57487106 |
| **SPATS2L** | Q9NUQ6 | 1.44826E-11 | 20.8269186 |
| **CXorf56** | Q9H5V9 | 1.79518E-11 | 22.04429579 |
| **ATAD3B** | Q5T9A4 | 2.50029E-11 | 24.08491898 |
| **PSMB3** | P49720 | 2.57109E-11 | 19.10311842 |
| **TFDP2** | Q14188 | 2.68949E-11 | 21.74063826 |
| **NARS2** | Q96I59 | 3.21775E-11 | 22.99020576 |
| **TFDP1** | Q14186 | 4.03805E-11 | 23.44895029 |
| **ARHGEF40** | Q8TER5 | 6.48736E-11 | 18.54688215 |
| **DST** | Q03001 | 6.67033E-11 | 23.83796215 |
| **DYNC1I2** | Q13409 | 1.16082E-10 | 18.75141859 |
| **ALOX12B** | O75342 | 1.22401E-10 | 19.90516663 |
| **CAMTA2** | O94983 | 1.51748E-10 | 18.84609842 |
| **TCEANC2** | Q96MN5 | 1.61163E-10 | 22.30599213 |
| **MICAL2** | O94851 | 1.91471E-10 | 22.78196573 |
| **LUC7L2** | Q9Y383 | 2.64243E-10 | 19.90857267 |
| **NALCN** | Q8IZF0 | 3.9284E-10 | 23.65266705 |
| **HSD17B12** | Q53GQ0 | 5.14354E-10 | 19.76246071 |
| **G3BP2** | Q9UN86 | 5.34248E-10 | 20.74985456 |
| **BMP1** | P13497 | 6.01063E-10 | 17.74047995 |
| **MCM5** | P33992 | 6.15221E-10 | 21.86290598 |

| | | | |
|---|---|---|---|
| **MTCL1** | Q9Y4B5 | 8.72964E-10 | 19.16291666 |
| **GTF2F1** | P35269 | 1.40001E-09 | 21.96182203 |
| **ARL6IP4** | Q66PJ3 | 2.82612E-09 | 20.32786083 |
| **ASB1** | Q9Y576 | 3.13137E-09 | 20.32534218 |
| **FHOD1** | Q9Y613 | 4.61619E-09 | 21.79634285 |
| **WRNIP1** | Q96S55 | 9.89487E-09 | 20.77000856 |
| **C1QBP** | Q07021 | 1.38315E-08 | 22.04387093 |
| **FARSB** | Q9NSD9 | 2.03539E-08 | 21.6620307 |
| **MCM3** | P25205 | 3.04929E-08 | 21.55954313 |
| **RAI14** | Q9P0K7 | 5.289E-08 | 22.09490585 |
| **TJP1** | Q07157 | 9.6431E-08 | 9.060780525 |
| **PRDX2** | P32119 | 9.48772E-07 | 2.160865784 |
| **VWA8** | A3KMH1 | 2.58649E-06 | 8.374697685 |
| **IGHG2** | P01859 | 2.87324E-06 | 2.267016888 |
| **RB1** | P06400 | 1.69138E-05 | 5.728475571 |
| **E2F3** | O00716 | 0.000123596 | 4.44920063 |
| **HS2ST1** | Q7LGA3 | 0.000805649 | 2.787140369 |
| **HDLBP** | Q00341 | 0.001226053 | 16.58484364 |
| **MYO1C** | O00159 | 0.00368581 | 1.961327553 |
| **IGHG1** | P0DOX5;P01857 | 0.004173774 | 1.447899342 |
| **C9orf43** | Q8TAL5 | 0.005461557 | 19.79761076 |
| **METTL15** | A6NJ78 | 0.006076881 | 18.81872368 |
| **SYNE2** | Q8WXH0 | 0.006637627 | 19.43278456 |
| **GTF3C1** | Q12789 | 0.007003875 | 15.98174143 |
| **GTF3C5** | Q9Y5Q8 | 0.008248568 | 17.46328402 |
| **MAP1A** | P78559 | 0.008712723 | 15.73902512 |
| **PUF60** | Q9UHX1 | 0.0089359 | 16.43379593 |
| **SLAIN2** | Q9P270 | 0.009466481 | 16.88302183 |
| **LRSAM1** | Q6UWE0 | 0.009942525 | 17.61318159 |
| **PLOD2** | O00469 | 0.009959096 | 15.37081671 |
| **CCAR1** | Q8IX12 | 0.010096252 | 16.58990526 |
| **GTPBP2** | Q9BX10 | 0.010515084 | 15.82120132 |
| **LCN1;LCN1P1** | P31025;Q5VSP4 | 0.010799184 | 16.08209801 |
| **SH3D19** | Q5HYK7 | 0.011342001 | 16.60372448 |
| **LOR** | P23490 | 0.011838207 | 15.43536043 |
| **FARSA** | Q9Y285 | 0.012004423 | 16.31003904 |
| **GTF3C6** | Q969F1 | 0.012004478 | 16.02385902 |
| **MYO9A** | B2RTY4 | 0.012006823 | 16.52427387 |
| **DDX39B;DDX39A** | Q13838;O00148 | 0.012006966 | 14.40969276 |
| **MGA** | Q8IWI9 | 0.012008256 | 15.91280937 |
| **PSMD13** | Q9UNM6 | 0.012008775 | 14.24829388 |
| **PSMB6** | P28072 | 0.012009292 | 16.03938675 |

| | | | |
|---|---|---|---|
| **SH2D3C** | Q8N5H7 | 0.012009504 | 15.85227156 |
| **RAB1A;RAB1B;RAB1C;RAB8B;RAB8A;RAB10;RAB13;RAB15** | P62820;Q9H0U4;Q92928;Q92930;P61006;P61026;P51153;P59190 | 0.012009763 | 16.35257006 |
| **SLC39A7** | Q92504 | 0.012013979 | 13.94072628 |
| **SRP9** | P49458 | 0.012014413 | 15.26320553 |
| **WDR1** | O75083 | 0.012015373 | 14.04478264 |
| **MANBA** | O00462 | 0.012015711 | 15.70327425 |
| **PSMA3** | P25788 | 0.012016117 | 15.26424742 |
| **PDIA6** | Q15084 | 0.012018752 | 16.17472506 |
| **TSEN15** | Q8WW01 | 0.012019648 | 15.62000895 |
| **INTS1** | Q8N201 | 0.012029182 | 16.50816965 |
| **ANKRD12** | Q6UB98 | 0.012034191 | 16.93458366 |
| **TMEM33** | P57088 | 0.012037901 | 16.90490389 |
| **LRP8** | Q14114 | 0.012045052 | 14.9916234 |
| **PTPRG** | P23470 | 0.012062199 | 17.77592945 |
| **PFKP** | Q01813 | 0.012066504 | 16.19368744 |
| **PSME3** | P61289 | 0.012072013 | 17.23930597 |
| **PRPF4B** | Q13523 | 0.012073328 | 17.01796865 |
| **QPCTL** | Q9NXS2 | 0.012084156 | 15.13104677 |
| **TSEN2** | Q8NCE0 | 0.01209207 | 16.30651617 |
| **SMC3** | Q9UQE7 | 0.01210472 | 14.55299091 |
| **SEPTIN9** | Q9UHD8 | 0.012117349 | 14.87176657 |
